# Computational assessment of blood flow heterogeneity in dialysis patients’ cardiac ventricles

**DOI:** 10.1101/301572

**Authors:** Sanjay R Kharche, Aaron So, Fabio Salerno, Ting-Yim Lee, Chris Ellis, Daniel Goldman, C W McIntyre

## Abstract

Dialysis prolongs life but augments cardiovascular mortality. Imaging data suggests that dialysis increases myocardial blood flow (BF) heterogeneity, but its causes remain poorly understood. A biophysical model of human coronary vasculature was used to explain the imaging observations, and highlight causes of coronary BF heterogeneity.

Post-dialysis CT images from patients under control, pharmacological stress (adenosine), therapy (cooled dialysate), and adenosine and cooled dialysate conditions were obtained. The data presented disparate phenotypes. To dissect vascular mechanisms, a 3D human coronary vasculature model was implemented. Simulations were performed to investigate the effects of altered aortic pressure and blood vessel diameters on myocardial BF heterogeneity which was quantified using relative dispersion, fractal dimension, and transmural BF distribution.

Imaging showed that stress and therapy potentially increased mean and total BF, while reducing heterogeneity. BF histograms of one patient showed multi-modality. Using the model, it was found that total coronary BF increased as coronary perfusion pressure (CPP) was increased. BF heterogeneity was differentially affected by large or small vessel blocking. BF heterogeneity was found to be inversely related to small blood vessel diameters. Simulation of large artery stenosis indicates that BF became heterogeneous (increase relative dispersion) and gave multi-modal histograms. The total transmural BF as well as transmural BF heterogeneity reduced due to large artery stenosis, generating large patches of very low BF regions downstream. Blocking of arteries at various orders showed that blocking larger arteries results in multi-modal BF histograms and large patches of low BF, whereas smaller artery blocking results in augmented relative dispersion and fractal dimension. Transmural heterogeneity was also affected. Finally, the effects of augmented aortic pressure in the presence of blood vessel blocking shows differential effects on BF heterogeneity as well as transmural BF.

Improved aortic blood pressure may lead to improved BF. Stress and therapy may be effective if they dilate small vessels. A potential cause for the observed complex BF distributions (multi-modal BF histograms) may indicate existing large vessel stenosis.

The intuitive BF heterogeneity methods used can be readily used in clinical studies. Further development of the model and methods will permit personalised assessment of patient BF status.

## 1. Introduction

Dialysis is a life prolonging treatment but significantly reduces quality of life due to its deleterious side effects on the heart. This mathematical modelling study explores some of the coronary vasculature based causes of myocardial blood flow (BF) heterogeneity.

### 1.1 Clinical motivation

The over 20% mortality among end stage renal disease (ESRD) patients receiving dialysis treatment is often due to cardiovascular complications (Collins et al., 2015). Dialysis is a repetitive sub-lethal ischemia that is known to produce cardiac contractile dysfunction as observed clinically (Breidthardt et al., 2012; Burton et al., 2009; Jefferies et al., 2011; McIntyre, 2010). ESRD patients also experience significantly reduced coronary blood flow (BF) (Dasselaar et al., 2009). The reduced BF may often occur due to calcification (McIntyre and Odudu, 2014; McIntyre et al., 2013) that reduces arterial diameters. Experimental observations have led to the belief that an increased resistance of blood vessels in the sub-endocardium promotes increased transmural BF heterogeneity (Algranati et al., 2011). Although non-invasive interventions may not affect existing large vessel structural defects such as stenosis, it is thought that adenosine stress and dialysate cooling therapy may improve myocardial BF by vasodilation of the smaller blood vessels. The interventions may also improve myocardial BF by improving aortic pressure. The mechanisms by which BF heterogeneity is affected remain unclear. Knowledge of the cause-effect relationships may permit design of future clinical trials and augment the precision of medications given to this critically ill group of patients.

### 1.2 Previous coronary vasculature theoretical models, extant experimental data

The predominantly dichotomous mammalian coronary architecture is complex with millions of arterial segments (Kassab et al., 1997a; Kassab et al., 1993). The larger left and right coronary arteries (∼ 3 mm diameters, 80 mm lengths) forming aortic ostia give rise to arterial trees, which together constitute the arterial vasculature. A very large number of pre-capillary arterioles deliver blood to capillary beds (∼ 0.06 mm diameters, 0.15 mm lengths). A spectrum of biophysical and anatomical properties of vasculature has been studied to permit patho-physiological investigations. The topology of coronary arterial trees has been quantified by Kassab et al. (Kassab et al., 1997a; Kassab et al., 1997b; Kassab et al., 1993) using silicon elastometer casts. In the absence of biophysical morphometry data, theoretical vasculature topologies can also be generated (Keelan et al., 2016). The topology data obtained from large animal hearts can be scaled to the human heart using clinical angiograms (Dodge et al., 1992). To permit generating a 3D geometry from the topology, the bifurcation properties of the network have been quantified. In accordance with Murray’s law (Murray, 1926a, b), the relationship between artery segment lengths, diameters, and bifurcation angles and planes has been described by Zamir and others (Zamir and Phipps, 1988; Zamir et al., 1983). Using the topology and Murray’s law, algorithms that distributed the topology as uniformly as possible in 3D space were developed. The algorithm, which may overall be called “space filling algorithm”, is based on the principles of self-avoidance and boundary avoidance and was developed by Beard and Bassingthwaighte (Beard and Bassingthwaighte, 2000). Several studies further developed the space filling algorithm which optimises vascular spatial distribution as well as delivery of BF to various parts of the heart (notably in Koimovitz et al. (Kaimovitz et al., 2010), Mittal et al. (Mittal et al., 2005), and Smith et al. (Smith et al., 2000)), and have either generated the complete or partial epicardial vasculature networks. Whereas vascular resistance is regulated by geometry alone, the BF and pressure at each location in the network also depends on the properties of fluid flowing through the network. Blood is a biphasic fluid and alterations of its viscosity, dependent on haematocrit, have been quantified by Pries et al. (Pries et al., 1996).The intricate problem of vasculature involves optimising relative arterial diameters, bifurcation angles, providing of BF to potentially empty regions, and being as widely distributed in the myocardium as possible. In the absence of experimental data it may be possible to generate virtual vasculatures based on biophysical principles and computational optimisation of cost functions (Kaimovitz et al., 2005; Karch et al., 1999; Keelan et al., 2016). The scientific problems arising in vasculature modelling based on available knowledge have been addressed in individual theoretical studies with particular aims. Alarcon et al. provide a design principle in light of complex blood rheology (Alarcon et al., 2005).

The effects of structural defects on haemodynamic distribution have been studied by Yang and Wang (Yang and Wang, 2013). The availability of mathematical-computational tools such as the 3D coronary vasculature models has encouraged the investigation of specific disease conditions in the heart (Zhang et al., 2014). Similar to the present study, a detailed model by Fung et al. (Fung et al., 2011; Fung et al., 2010) has been developed to assist in evaluation of imaging hearts with perfusion defects. An extension of previous and presented models will incorporate the multi-scale nature of delivery of oxygen (Goldman and Zahra Farid, 2017; Mason McClatchey et al., 2017) to myocardial tissue. Although several studies have characterised the properties of the coronary vasculature, the use of this vast basic science knowledge for clinical purposes remains acutely limited.

In this study, we endeavoured to exploit organ level vasculature modelling to investigate potential causes for our clinical myocardial BF heterogeneity observations.

## 2. Methods

### 2. 1 Clinical imaging

It is thought that peritoneal dialysis increases coronary BF heterogeneity in patients. CT imaging was performed to test whether adenosine, cooled dialysate, and adenosine combined with cooled dialysate can enhance myocardial BF, and reduce BF heterogeneity.

#### 2.1.1 Patient recruitment

Three chronic end stage renal failure patients, aged between 58-63 years, were recruited from the London Health Sciences Centre Peritoneal Dialysis Program (London Ontario, Canada). Each patient had been on peritoneal dialysis for a minimum of 3 months prior to recruitment. All patients were informed regarding the study, after which written consents were obtained in accordance with the hospital and university procedures. All participants provided informed and written consent. The study protocol was approved by the research ethics board at Western University (London, Ontario, Canada).

#### 2.1.2 Imaging study protocol

Briefly, each patient was scanned four times in two study visits. During the first visit, a glucose based peritoneal dialysis (according to their prescription) was administered at a physiological temperature of 37°C, after which they were scanned with and without adenosine stress. During the second visit, patients were administered the peritoneal dialysis dose but with a cooled dialysate (32.5°C), after which they were scanned with and without adenosine stress.

To assist the dynamic contrast enhanced CT scanning, a contrast agent (lopamidol) was administered. The heart rate was also reduced using a beta-blocker that permitted a longer diastolic phase in the left ventricle. The details of the imaging protocol and image processing that computed the BF maps are given in **Supplementary Methods Section S1**.

#### 2.1.3 Patient blood pressure

The diastolic blood pressures values ranged between diastolic 71 mmHg to 85 mmHg, and systolic values between 130 mmHg to 210 mmHg. Other clinical laboratory measurements were not considered in this study.

#### 2.1.5 3D heart segmentation

The 2D registered BF map slices were segmented by a physician semi-automatically using Fiji/ImageJ (Schindelin et al., 2012). Each slice was first segmented to remove non-myocardial tissue signals. Subsequently, the image was thresholded to between 0 and 600 ml/mg/min to selectively remove residual signals pertaining to intracardiac (left and right ventricle chambers) blood, while preserving signals providing coronary BF distribution. The slices were stacked to reconstruct each patients BF maps under each of the four clinical conditions (see sec 2.1.2). The reconstruction was stored in a structured array of 0.5 (X) x 0.5 (Y) x 5 (Z) mm^3^ array. The structured array representation was used to quantify BF heterogeneity (see below).

### 2.2 Model construction

#### 2.2.1 Human ventricle anatomy to contain vasculature geometry

A representation of the human ventricles was constructed to permit generation of vascular geometry within it. An idealised representation bound by truncated ellipsoids was constructed (Göktepe and Kuhl, 2010). The dimensions of the anatomy are detailed in **Supplementary Methods, Section S2 and Figure S1**. The coronary vasculature geometry was generated within the ventricular anatomy.

#### 2.2.2 Topology of coronary vasculature based on morphometry biophysical data

The porcine coronary vasculature morphometry (Kassab et al., 1993) was used to construct arterial tree topology. The stochastic morphometry data consists of arterial segment connectivity matrices, segment lengths, and segment radii. In this study, coronary arterial topologies were constructed as binary trees, one each for the right and left ventricle. The segments, defined as parts of arteries between two consecutive bifurcation nodes, in the trees were numbered according to the Strahler number (SN) ordering (Strahler, 1957). Within this numbering system the largest arteries are a series of segments of the same SN, namely the left anterior descending (LAD) and the right coronary artery (RCA), have a SN of 11. The left circumflex artery (LCX) which has a SN of 10 emerges at a bifurcation of the LAD, and provides BF to a sub-tree in the left ventricle. Whereas the larger vessels (SN 9 to SN 11) are restricted to the epicardial surface, smaller vessels (SN 6 to SN 8) provide BF transmurally (Kaimovitz et al., 2005). As a computationally manageable approximation that permitted simulation of whole heart BF, the arterial trees were generated for SN 6 to SN 11, where SN 6 was identified based on its diameter and the number of bifurcations that would be needed to reach the capillary level (SN 0). First, arterial elements, defined as a series of connected segments of the same SN, were generated stochastically using the segment to element ratios in the morphometry data (**Supplementary Methods Section S3**). The elements were then assembled, again stochastically, using connectivity matrices (Supplementary Methods, Section S3) (Kassab et al., 1993) to stochastically generate multiple instances of the whole heart’s coronary arterial binary tree topologies. Each element was assigned a constant radius along its length adapted from the experimental data of Kassab et al. in accordance with previous modelling studies (Beard and Bassingthwaighte, 2000; Smith et al., 2000). Finally, the segments were assigned lengths stochastically (**Supplementary Materials Section S3**) (Kassab et al., 1993). The total tree lengths were bounded to avoid non-physiologically short or long trees (Kaimovitz et al., 2005). The total tree length of RCA was limited to between 120 mm and 192 mm, and those of LAD and LCX were both limited to between 100 mm and 160 mm.

Using the segment lengths (L) and radius (r) information assigned during topology generation, and using a blood viscosity value of μ =3.6 × 10^−3^ Pa (Keelan et al., 2016), a value of resistance to flow (R) in each segment was assigned using the relationship:

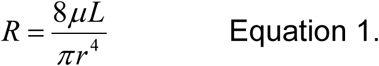

The downstream resistance at any bifurcation node was then computed recursively. Elements that include segments of SN 8 to SN 11 constitute epicardial vessels (Kaimovitz et al., 2005; Kassab et al., 1993; Smith et al., 2000).

To permit generation of arterial geometry, pressure boundary conditions were imposed on the inlets and terminal nodes of the arterial tree topologies and BF in each segment and pressure at each internal node were computed recursively. At the terminal nodes (SN 6), the pressure was set at 20 mmHg. At the inlets, the pressure was set at 100 mmHg under control conditions, and varied to simulate disease, stress, or therapy. The boundary conditions, conservation of flow at bifurcation nodes, and Poiseuille’s law for steady state flow, ΔP (pressure) = R (resistance) x Q (blood flow) in each segment and in whole trees, permitted calculation of pressure at each bifurcation node, and BF through each segment of the arterial trees.

#### 2.2.3 Generation of arterial tree geometry ensemble

The roots of the RCA and LAD arterial trees were placed approximately at their aortic ostia locations (Kaimovitz et al., 2005; Keelan et al., 2016; Smith et al., 2000). The largest arteries were placed either along the right or left epicardial surfaces (RCA and LCX respectively), or traversing from the base (AV border) to the apex (LAD). Doing so provided a starting point for the self-avoidance and boundary-avoidance algorithms as described by Beard and Bassingthwaighte (Beard and Bassingthwaighte, 2000). The methods are described briefly for completeness.

To implement the self-avoidance algorithm, a vascular supply vector specific to an arterial node, *v_s_*, was computed as follows:

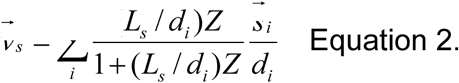

where *s⃗_l_* − *x_c_* − *x_i_ S_i_* = *x_c_* − *x_i_*, *x_i_* is upstream position of all segments of same or higher order with coordinates, *x_c_* is the upstream position of the current generating segment; *d_i_* is the magnitude of the vector *s⃗_i_*.. *Z* is the avoidance exponent and was taken to be 2 throughout the arterial trees (Beard and Bassingthwaighte, 2000).

In addition to self-avoidance, boundary avoidance was implemented to constrain the vascular trees within the myocardium predefined by the ellipsoidal ventricular epicardial and endocardial walls. The unit normal vectors required to implement self-avoidance were computed as outward (endocardial) or inward (epicardial) normals, *n_j_*, to the ellipsoidal surfaces defining the bi-ventricular wall boundaries of the myocardial geometry (**Supplementary Methods, Section S2**). Using the unit normals and the supply vector, another vector (*D*) directed away from the previously defined arterial nodes as well as from the boundaries was defined as:

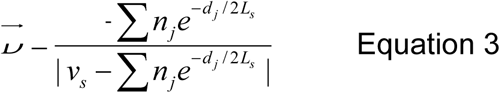

Further, the combination of *v_s_* and *D* was used to construct initial positions of the left and right segment distal nodes, as described in recent studies (Fung et al., 2011; Tamaddon et al., 2016; Yang and Wang, 2013). *v_s_* and *D* were combined to construct a combined branching vector *v_d_* as follows:

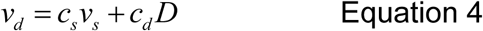

where *c_s_* and *c_d_* were both set to 0.5. The branching plane, in terms of it’s vector normal, was computed using the combined branching vector (Eq. 4) and the parent segments direction (Fung et al., 2011). Branching angles ϴ_L_ and ϴ_R_ for the two daughter branches were computed using the estimated BF in each of the daughter segments (Fung et al., 2011; Hacking et al., 1996; Zamir, 1976). Using the branching angles, the combined branching vector was rotated (see **Supplementary Methods Section S5**) by ϴ_L_ and - ϴ_R_, or - ϴ_L_ and ϴ_R_ to construct the directions along with the daughter vessel nodes were to be placed. These vectors were then used to define the positions of the two child nodes, x_dL_ and x_dR_, emerging from the bifurcation as

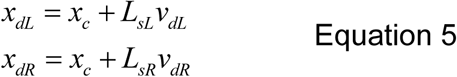

In case the computed locations (Equation 5) were not in within the geometry, they were then incrementally moved away from the violating boundary till *x_d1_* and *x_d2_* fell within the tissue. The incremental movement of the nodes was an iterative process and was performed for 200 iterations. sub-trees that were not assigned positions were then pruned. Upon assignment of positions to all segments in the trees, the pressure and BF were computed again using Poiseuille’s law.

As the arterial trees constructed in this study were limited to SN 6-11, an ensemble of 540 instances of the coronary vasculature (consisting of RCA, LAD, and LCX sub-trees) were constructed to allow accurate estimation of BF in simulation experiments.

#### 2.2.4 Construction of BF map in 3D, quantification of heterogeneity

##### BF map construction

The model was divided into 1 mm^3^ (high resolution) or 2 mm^3^ (low resolution) voxels. BF to a given voxel contained within the ventricular walls was assigned as the sum of BF received through all SN 6 terminals that had coordinates belonging to the volume of that voxel. Such a voxelised distribution of BF was computed for each instance in the ensemble. An average over the complete ensemble was performed to give a BF map. This BF map was used in computing measures for heterogeneity.

##### BF histograms

The ranges of BF values in the voxelised BF maps were binned, or grouped, into 600 bins for the imaging data, and 100 bins for the modelling data. The numbers of values in each bin were counted and a BF histogram was constructed. The means and standard deviations of the BF histograms were computed.

##### Probability distribution function of relative BF, fractal dimension

Probability distribution functions (PDFs) of relative flow were calculated to permit estimation of relative dispersion (Bassingthwaighte et al., 1989). To do so, the voxelised BF values were first normalised as

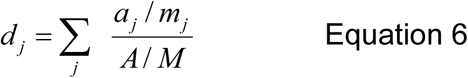

Where *a_j_* is the perfusion in the piece, *m_j_* is the mass of the piece, *A* is total perfusion in the 3D BF map or the imaging data, and M is the total mass of the ventricles. The masses of 1 voxel in the imaging data, and 1 mm^3^ voxel in the modelling data were assumed to be 1. The probability density of each bin was computed to give a probability density function (PDF) over a finite interval histogram. The area under this histogram and its mean were confirmed to be unity, to ensure that this histogram represented a PDF. The standard deviation of this PDF was taken to be the relative dispersion, RD. RD was computed at two resolutions and the lower resolution was taken to be the reference resolution. Fractal dimension, *D*, was computed using these two values of RD using the relationship (Bassingthwaighte et al., 1989):

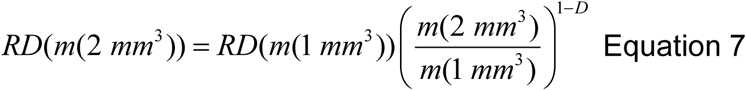

##### Transmural BF heterogeneity

To characterise transmural BF heterogeneity, the ventricles were divided into slices of 1 mm thickness that were equidistant from endocardial surfaces. Starting with the structured grid of the BF map, the shortest distance of each 1 mm^3^ voxels mid-point from the endocardial surfaces was computed. As the surfaces are idealised truncated ellipsoids, a geometric method detailed in **Supplementary Section S4** was implemented to generate distance maps. Using the distances, each voxel’s BF was assigned to a 1 mm thick layer. In particular, the voxels with distances between 1 and 2 mm from the endocardial surface were considered as the sub-endocardial layer.

#### 2.2.5 Simulation experiments

After characterisation of the baseline, or control, behaviour of the model, simulation experiments were performed. To simulate the effect of adenosine or dialysate cooling, the boundary conditions in terms of inlet pressures were varied from low (30 mmHg) to high (200 mmHg), and BF distribution calculated for each value of inlet pressure. To simulate the effect of dialysis or pre-existing structural conditions, vessels were constricted (stenosis). Stenosis of vessels consisted of either blocking the largest vessels, blocking of vessels that had a particular SN, or other structural characteristics. Finally, the combined effects of altered inlet pressure and blood vessel blocking were simulated.

#### 2.2.6 Numerical methods

The topology-geometry algorithms were implemented as a serial computer program using in house developed codes in C language, and ran on national High Performance Computing Services provided by Compute Canada. Simulation experiments and data analysis were performed using a local cluster. In both cases, the large number of simulations were optimally performed using serial farming job arrays which exploited the GNU parallel LINUX/UNIX utility (Tange, 2011).

## 3. Results

### 3.1 CT imaging based BF heterogeneity in patient ventricles

A representative 3D myocardial BF map which was reconstructed from thoracic CT images is shown in Figure 1, Row 1 and BF histograms in Figure 1, Row 2. The BF histograms were smoothed using Bezier smoothing and normalised to their respective maximum values to highlight the shifts in peaks. In comparison to control (red lines), adenosine (cyan lines) shifted the histogram peak to higher values. Dialysate cooling (green lines) shifted the histogram’s peak in an inconclusive manner. Adenosine with dialysate cooling (blue lines) shifted the peaks to higher values. In case of patients 1 and 2, the histograms are unimodal whereas in case of patient 3, all histograms are bi-modal with one major peak at a high BF value, along with a secondary peak at a low BF value. The BF histograms were converted to PDFs of relative flow (see Methods) (Figure 1, row 3). The standard deviations of these PDFs provided relative dispersions (RDs) at 1 voxel resolution. A similar PDF at a 4 voxel resolution was used to compute fractal dimensions (FDs) (Figure 1, Row 4). In case of all patients, adenosine reduced the 1 voxel RD and FD indicating a BF heterogeneity reducing effect. FD as regards dialysate cooling was either reduced in patients 1 and 2, or increased in case of patient 3. We noted that the BF histograms and PDFs for patient 3 are bi-modal.

**Figure 1.**
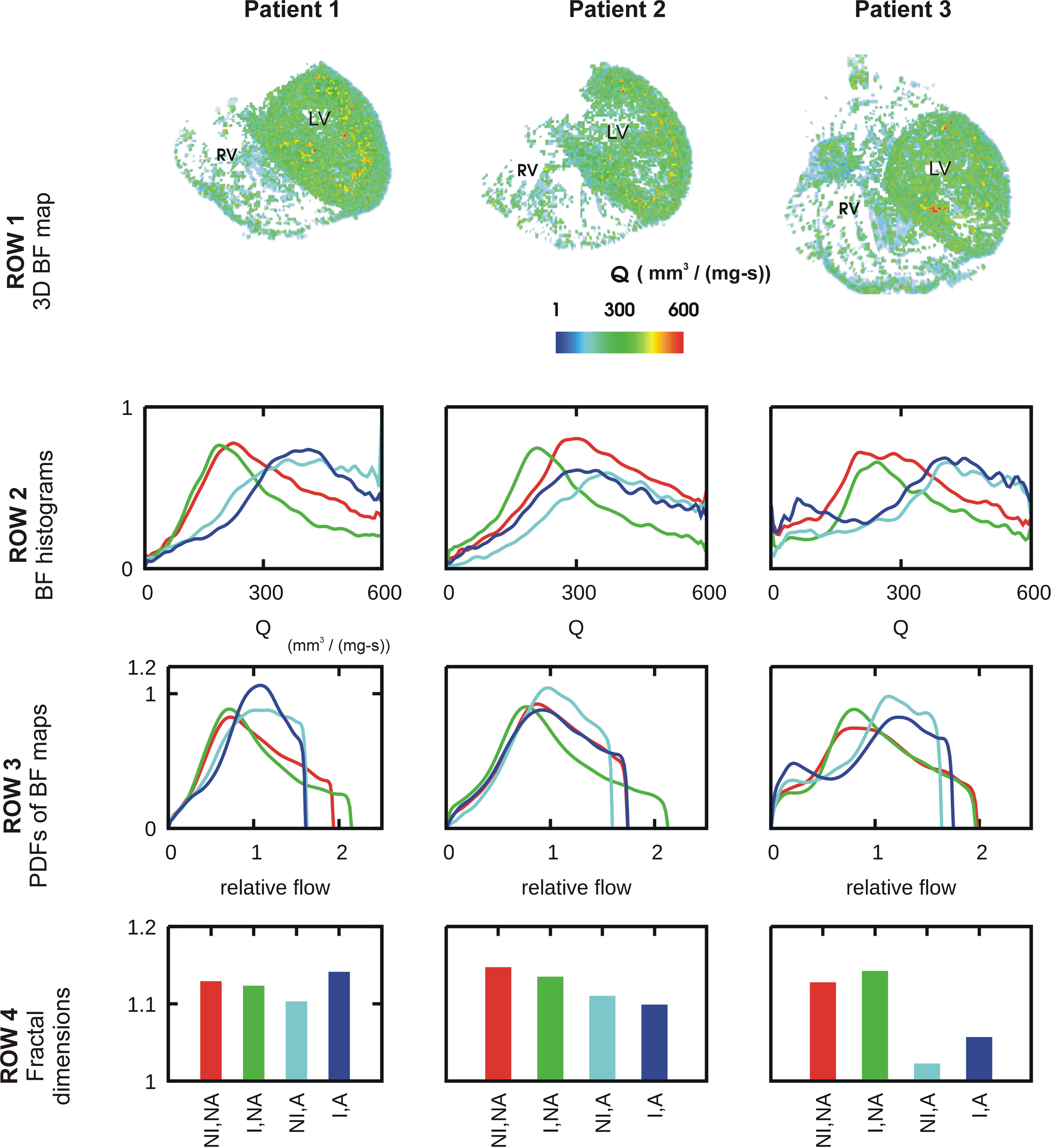
Characteristics of BF heterogeneity in three patients. Row 1: Representative 3D reconstruction of a myocardial BF map segmented from CT images. In the BF map instance (Ai), BF is shown in each segment. In the BF distributions (Aii), the average BF delivered by terminal segments to voxels is shown. In rows 2-4, data for control (red), cooled dialysate (green), adenosine treated (cyan), and cooled dialysate with adenosine treated (blue) are shown. Row 2: Histograms of BFs in the 3D reconstructions. Row 3: PDFs of relative BF computed from corresponding histograms of row 2. Row 4: FDs computed using the corresponding PDFs.

### 3.2 Modelling results

#### 3.2.1 Control in silico BF map and transmural heterogeneity

The control *in silico* BF map and its properties are illustrated in Figure 2. An instance of BF distribution in one of the 541 instances is shown in Figure 2, Ai. The average of 541 instances, voxelised at resolution of 1 mm^3^, was used to represent the control BF map (Figure 2, Aii). Histogram of BF, based on the BF map, shows that the mean 1 mm^3^ resolution BF was approximately 3.7 mm^3^/s. Histograms at 1 mm^3^ and 2 mm^3^ resolutions were used to construct the PDF of relative flow (Figure 2, Bii shows the 1 mm^3^ resolution PDF). The FD was found to be 1.18 showing a relatively low BF heterogeneity in the control BF map. The control *in silico* BF maps approximate error analysis was performed. FD was computed using an increasing number of instances. The values of FD were found to fit an exponential decay curve exactly (Figure 2, Biii). From the fitted curve, it was found that the asymptotic value was 1.14. The difference between our control BF map and the asymptotic value was 7%, and therefore the error was deemed to be negligible.

**FIGURE 2.**
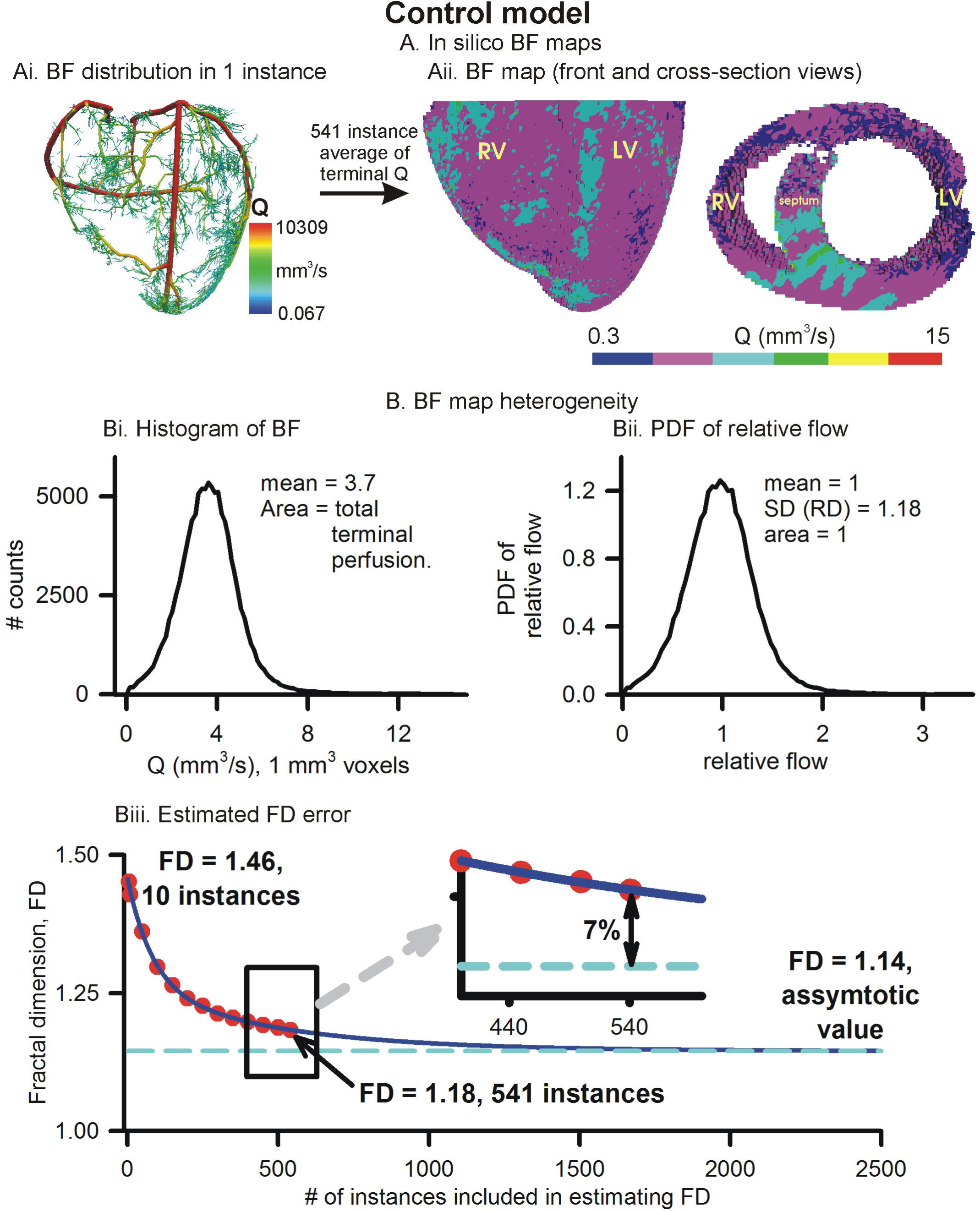
Simulated BF map and quantification of BF heterogeneity. Ai. An instance of vasculature structure showing BF distribution. Aii. Front and crosssectional views of simulated BF maps using the full 541 instances ensemble. Bi. Histogram of simulated BF map. Bii. PDF of relative perfusion which provides the relative dispersion (RD), also termed fractal dimension (FD). Biii. Difference between FD using complete ensemble (1.18) and estimated asymptotic value (1.14).

Further, the transmural BF heterogeneity was quantified (Figure 3). BF within layers of 1 mm thickness, computed from the distances of each 1 mm^3^ voxel from the endocardial surfaces (Figure 3, Ai), in the right and left ventricles are shown in Figure 3, Bi and Bii respectively. The epicardial layers in the right ventricle received significantly lower BF than in the left ventricle. The endocardial layers received a significant BF. In both the right and left ventricles, it was found that BF had a minimum in the sub-endocardial layers (Figure 3, Bi and Bii, gray boxes).

**FIGURE 3.**
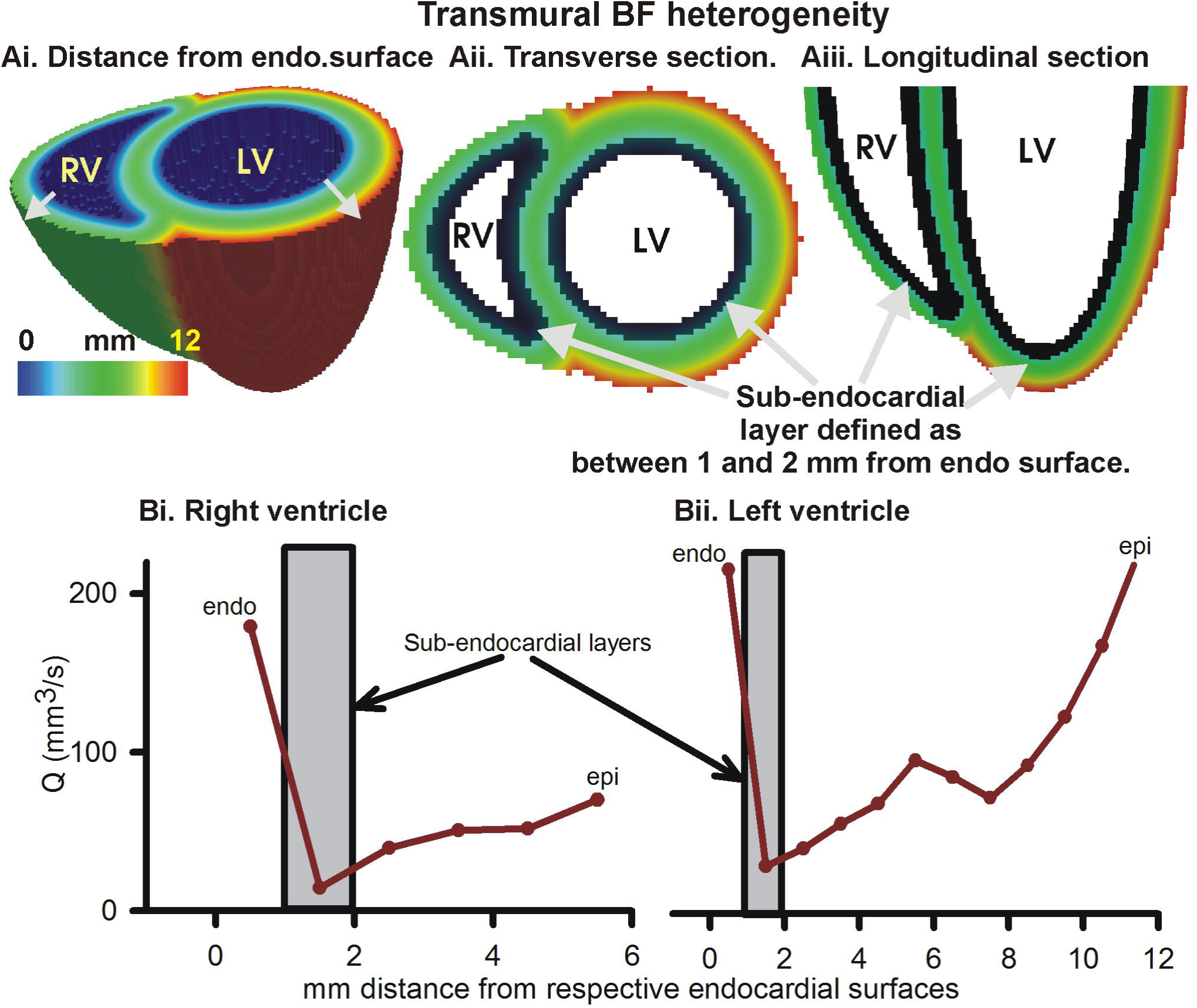
Transmural BF heterogeneity in the simulated BF map. Ai. Distance of each point in left and right ventricular tissue from respective endocardial surfaces. Increasing distance is shown by gray arrows. Aii. Cross-section at the base showing distance from endocardial surface. Aiii. Longitudinal section showing distance from endocardial layer. Black layer in Aii and Aiii shows sub-endocardial layer. Bi. Measured total perfusion in each 1 mm thick layer of the model right ventricle. Bii. Measured total perfusion in each 1 mm thick layer of the model left ventricle. Perfusion in sub-endocardial layer of RV and LV is shown in the gray boxes.

#### 3.2.2 Improved aortic pressure improved overall BF, heterogeneity unaltered

BF distributions under varying inlet pressure boundary conditions were simulated (Figure 4). Figure 4, A shows three representative simulated BF maps at low (30 mmHg), control (100 mmHg), and high (200 mmHg) inlet pressures, representing aortic pressures. The BF histograms are shown in Figure 4, B. At low pressure (sub-physiological to clearly show the effect), the histogram shows total BF to be low (area under curve of BF histogram). As inlet pressure was increased, the total BF also increased. The linear relationship between total BF and aortic pressure in Figure 4, C shows the constant resistance of the unaltered vascular structure. PDFs of relative perfusion (Figure 4, D) confirmed that the RD as well as the consequent FD remained unchanged, due to unaltered underlying coronary structure. The transmural BF was also unaltered (**Supplementary Figure S3**).

**Figure 4.**
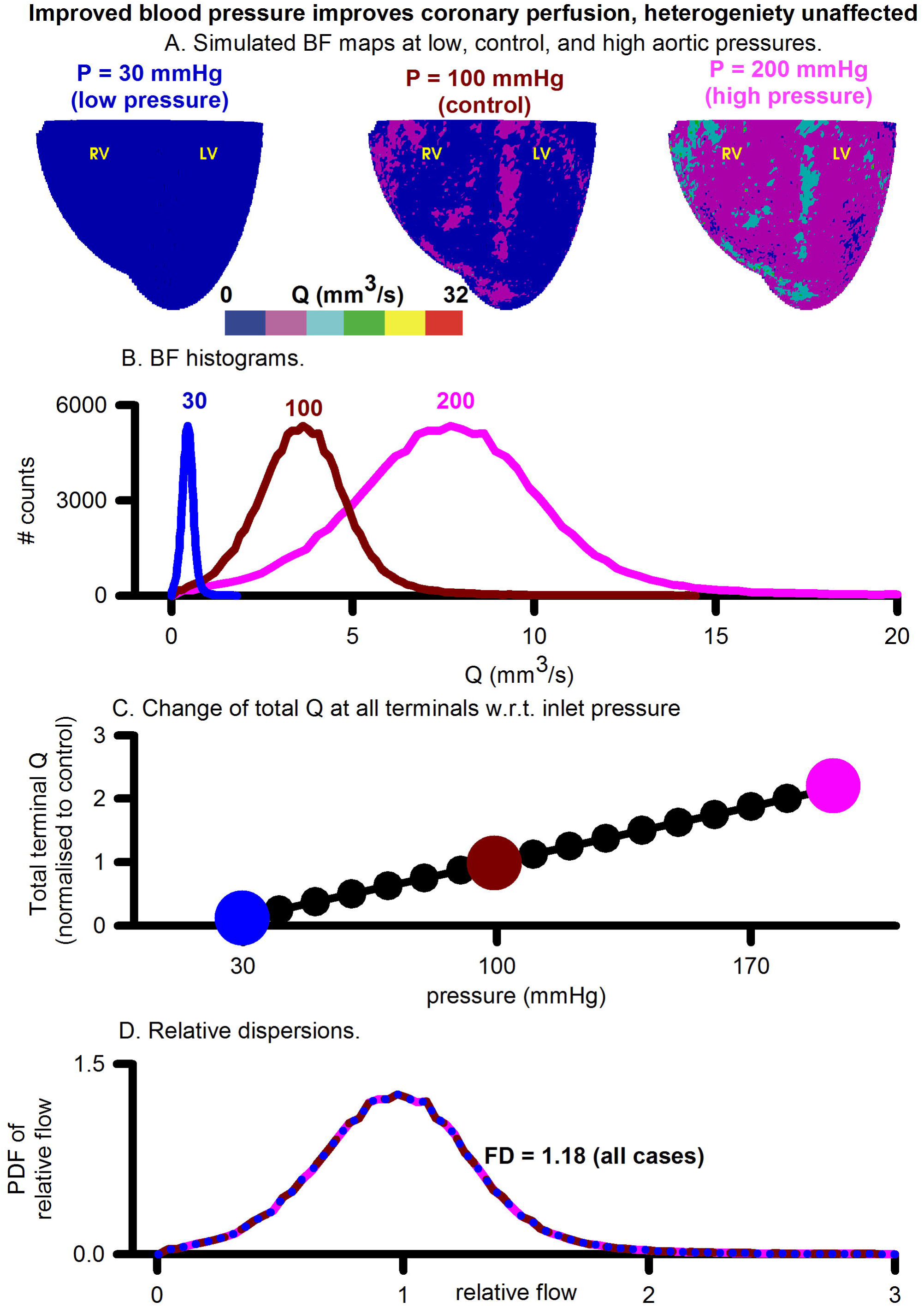
Assessment of the effect of increasing aortic (inlets) pressure. A. Simulated BF maps at 30 mmHg (left), 100 mmHg (center) and 200 mmHg (right). B. BF histograms at 30 mmHg (blue), 100 mmHg (red) and 200 mmHg (pink) showing increase of total perfusion with increase of inlet pressure. C. Total BF at terminals as a function of inlet pressure. Symbols show pressure values where simulations were performed. Large coloured circles show the 30 mmHg, 100 mmHg, and 200 mmHg values. D. Colour coded PDFs of relative perfusion, which were found to be identical for all pressure values.

#### 3.2.3 Large artery blocking (stenosis) promotes BF bi-modality, generates low BF distal regions

The effects of blocking an arbitrarily chosen LAD segment are illustrated in Figure 5. Pressure distributions and BF maps under control (r = 1.53 mm, Figure 5, Ai) and severe stenosis (r = 0.01 mm, Figure 5, Aii) illustrate the emergence of low BF distal regions, distal to the severe stenosis. At mild stenosis (r ≥ 0.25 mm), the PDFs of relative BF were found to be unimodal (Figure 5, Bi). In contrast, at severe stenosis (r < 0.25 mm), the histograms became bimodal with two distinct peaks, one each at low and high BF values (Figure 5, Bi). As the severity of stenosis was increased, the FD was found to progressively reduce from 1.18 (control) to 1.01 (severe stenosis) (**Supplementary Figure S4**). Severe stenosis also reduced the total perfusion by approximately 15% (**Supplementary Figure S4**). However, increased BF heterogeneity is reflected in the increase of RD as the severity of stenosis was increased (Figure 5, Bii). The region of BF distal to the location of stenosis experienced a progressively reduced pressure gradient as shown in Figure 5, C. As the stenosis segment was in the LAD and due to the approximate nature of the model (terminals at SN 6), the right ventricle transmural heterogeneity was unaffected (Figure 5, Di). However, the amount of BF received by all layers in the left ventricle reduced progressively as severity of stenosis was increased. Further, the transmural heterogeneity was found to reduce as radius of chosen LAD segment was reduced (Figure 5, Dii).

**Figure 5.**
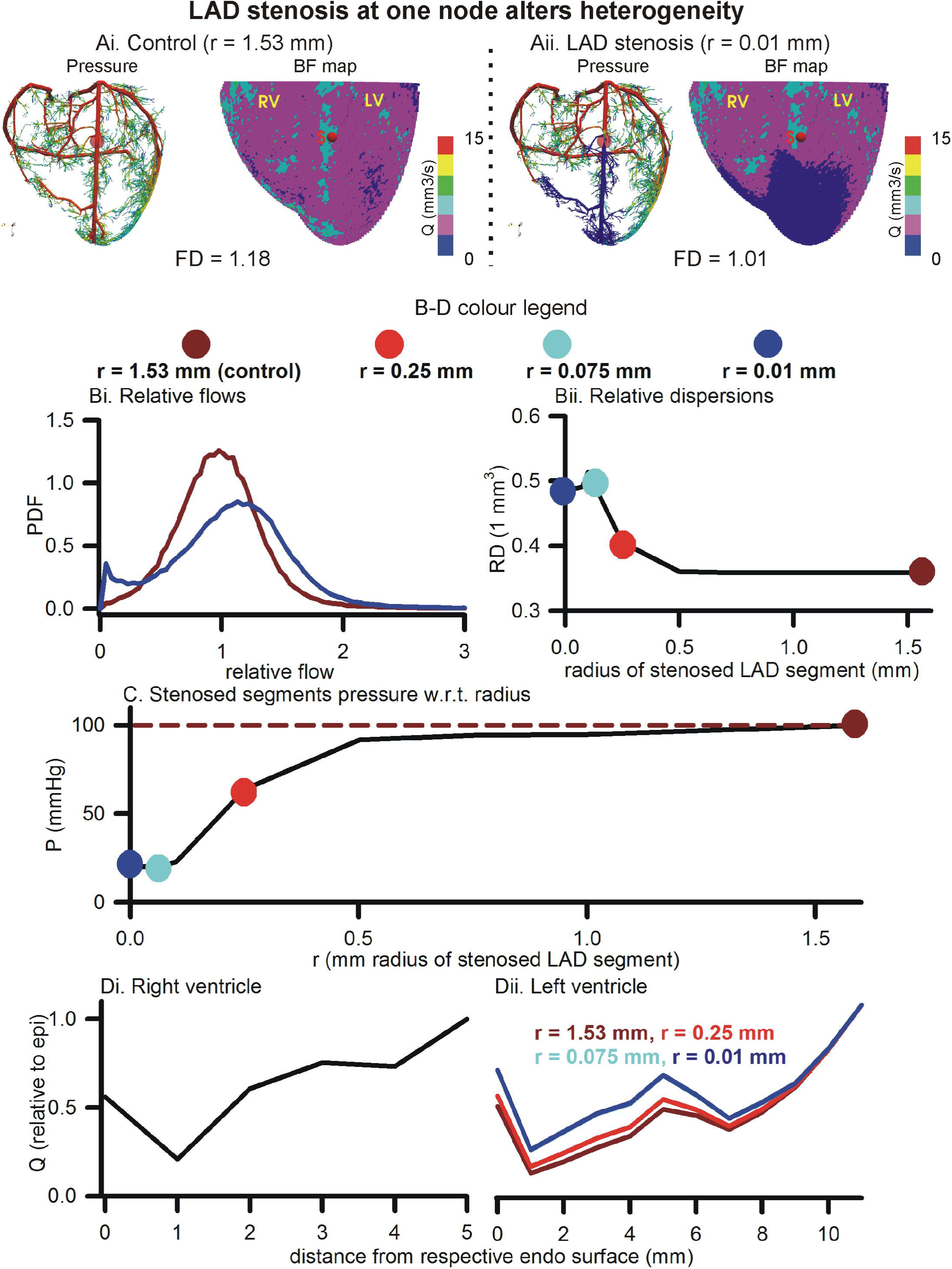
Effect of stenosis in large artery segment. Ai. Control pressure distribution in vasculature (left) and BF map (right). The control radius of the arbitrarily chosen LAD segment was r = 1.53 mm. Aii. Pressure (left) and BF map (right) under severe stenosis at the chosen LAD segment was r = 0.01 mm. In B-D, the colour coding is used to represent segment radii, r = 1.53 mm (dark red), r = 0.25 mm (red), r = 0.075 mm (cyan), and r = 0.01 mm (blue). Bi. PDFs of relative perfusion. Bii. Relationship between RD and stenosed segment radius. C. Relationship between pressure at stenosed segment and its radius. Di, Dii. Transmural BF heterogeneity in right (left) and left (right) ventricles.

#### 3.2.4 Blocking all large arteries (stenosis) promotes BF bi-modality, blocking smaller arterioles increase BF heterogeneity

The roots of arterial sub-trees of a given SN order (SN 6 to 10) were blocked by 90% of their control radius to generate BF maps (Figure 6, Ai). Blocking of root segments of SN 7 to SN 10 sub-trees altered the BF patterns. Blocking of terminals (SN 6) caused an apparent overall reduction of BF. Instances of the corresponding pressure distributions are shown in Figure 6, Aii. The BF maps and pressure gradients indicate that the volume of tissue affected is related to the SN order of the arterial sub-tree that was blocked. Relative dispersion of PDFs (Figure 6, B) increased (RD = 1.22, also see **Supplementary Figure S5**) when SN 6 terminals were blocked. The PDF was seen to be uni-modal. Blocking root segments of SN 7 or higher sub-trees also increased relative dispersion, but also gave rise to bi-modal PDFs with peaks at low and high values (Figure 6, B). The amplitude of the lower relative flow peak was greater at SN 10 blocking as compared to SN 7 blocking. The total BF when each SN order was blocked is shown in Figure 6, C. Blocking of the smallest arteries had the most significant effect of reducing total coronary BF. Blocking of SN 7-9 arteries had a relatively less impact. Blocking SN 10 arteries also reduced total coronary BF significantly. Blocking of arteries also affected transmural heterogeneity (Figure 6, D). In the right ventricle, blocking SN 7 maximally increased transmural heterogeneity. On the other hand, blocking SN 6, or SN 8-10 reduced transmural heterogeneity. In the left ventricle, blocking SN 6 or SN 10 increased transmural heterogeneity maximally. Blocking of SN 7, 8, or 9 reduced heterogeneity.

**Figure 6.**
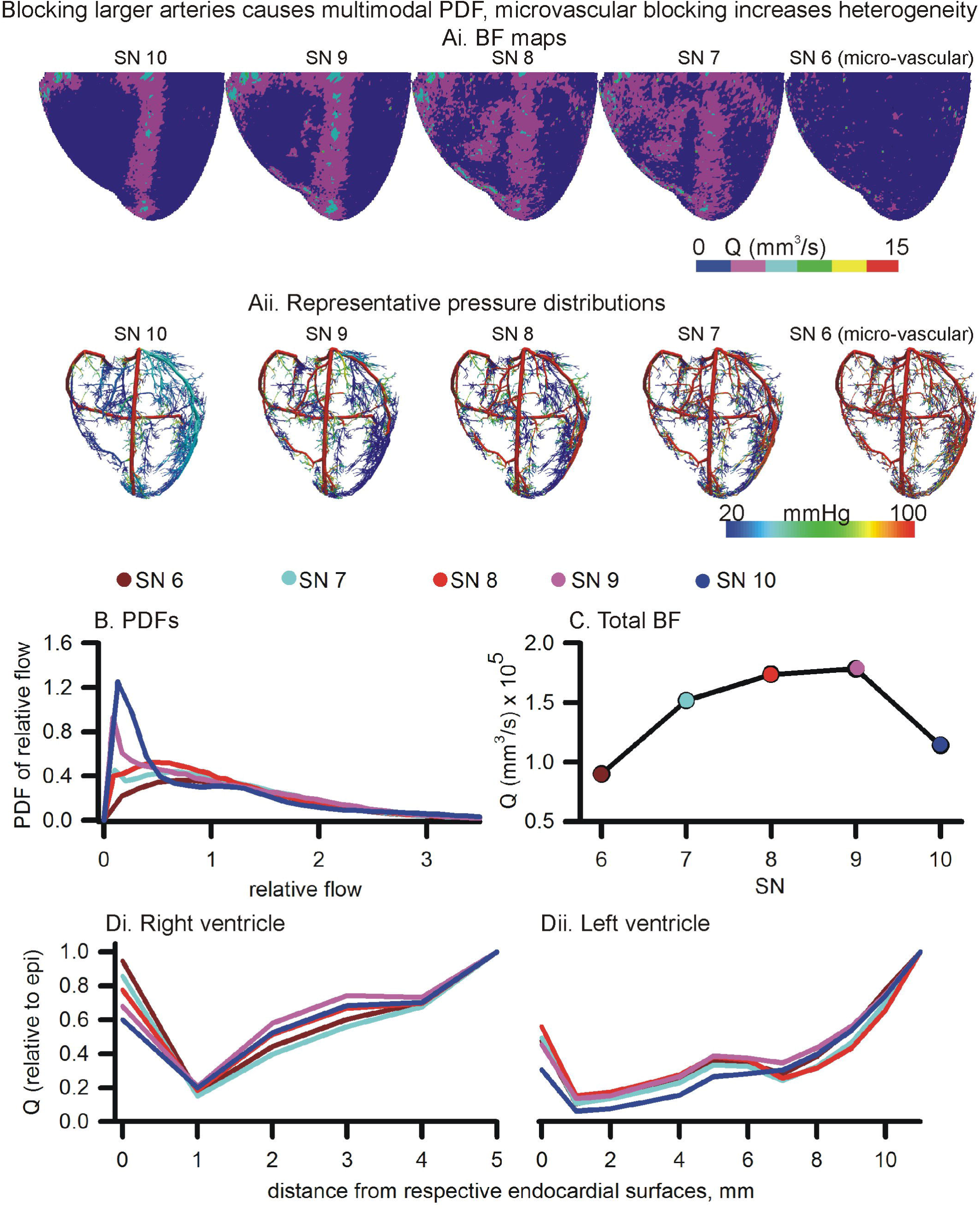
Effects of blocking arteries at given order on perfusion and heterogeneity. Ai and Aii. BF maps (Ai) and pressure distributions (Aii) when arteries of various orders were blocked. B. PDFs of BF maps. C. Total BF at terminals when that order arteries were blocked. Di, Dii. Epi-endo heterogeneity alterations under blocking of various orders of arteries.

#### 3.2.5 Altered inlet perfusion in the presence of blocked arteries

Simulation of diseased, or adenosine and dialysate cooling therapy induced alteration of aortic pressure in the presence of coronary structural defects are illustrated in Figure 7. Blocking of SN 6 terminals (70, 80, or 90 % of control radius values) increased the FD (Figure 7, A). In contrast, blocking of SN 10 sub-trees reduced the FD. Transmural heterogeneity was unaffected by blocking of SN 6 terminals (Figure 7, B). However, as the severity of SN 10 blocking increased, the transmural heterogeneity was found to be increased. Blocking SN 6 vessels has a greater impact on reducing total BF in comparison to blocking SN 10 vessels (Figure 7, C). BF reduced as aortic pressure reduced.

**Figure 7.**
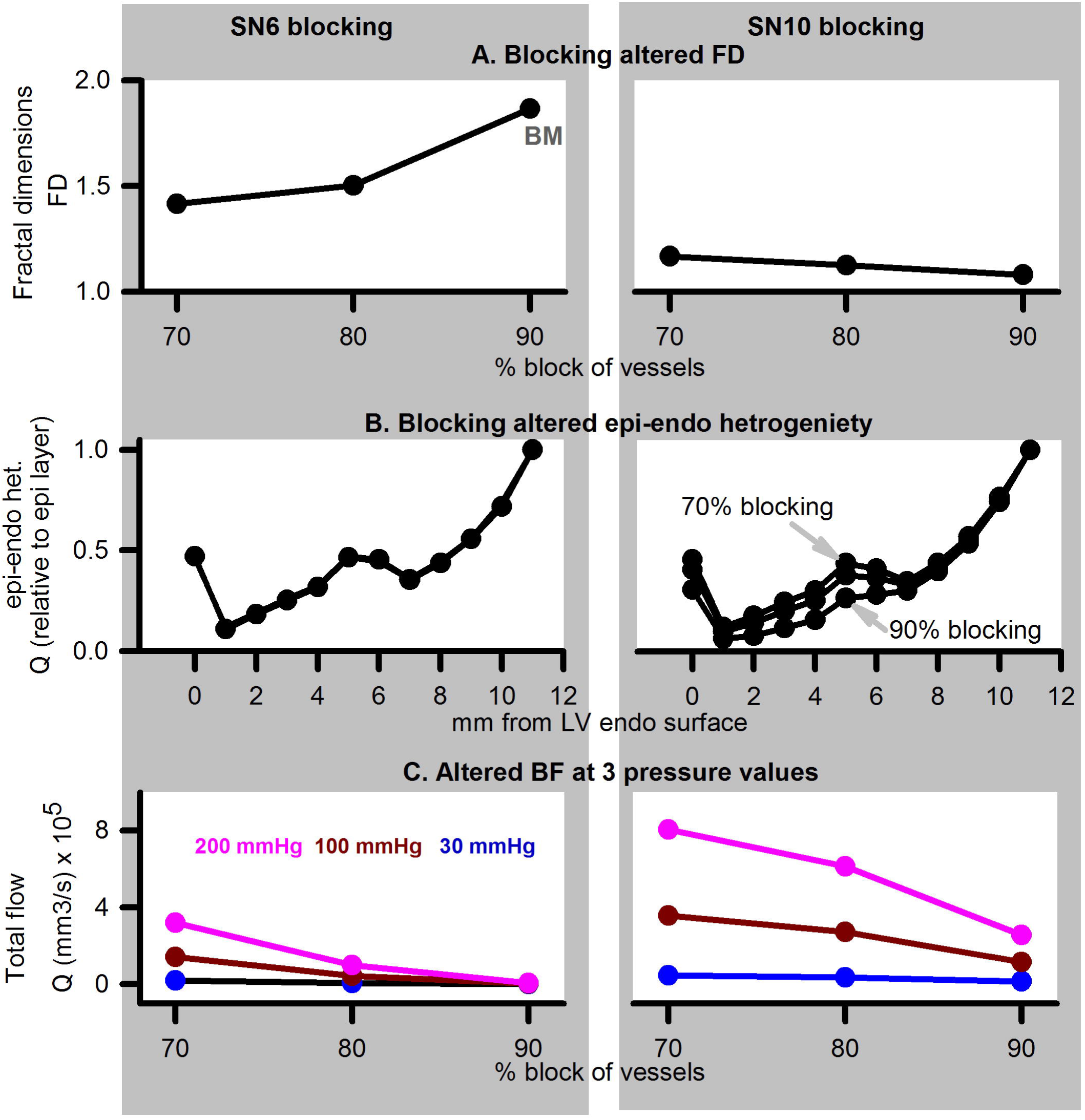
Simulation of disease (reduced pressure) or adenosine and dialysate cooling (therapy) in the presence of structural defects. Left column shows data for SN 6 blocking, while right column shows data for SN 10 blocking. A: FD when vessels were blocked by 70, 80, or 90% of their control values B: Transmural BF heterogeneity. C: Total BF when vessels were blocked by 70, 80, or 90% when applied inlet pressure was 200 mmHg (pink), 100 mmHg (dark red), or 30 mmHg (blue).

## 4. Conclusions and Discussion

The main conclusions of this study are:

a. Clinical imaging can provide information regarding BF alterations in patient ventricles. Total BF can become increased due to pharmacological (adenosine) and therapeutic (dialysate cooling) interventions. Although BF heterogeneity is also potentially affected by the interventions, the information is confounded by underlying vascular structural disease such as large vessel stenosis.
b. Computer modelling indicates that total coronary BF is increased by improved aortic pressure, and also by vasodilation of coronary blood vessels.
c. The causes of BF heterogeneity appear to be multi-fold. Small vessel constriction promoted increase of dispersion but maintained uni-modality of BF histograms. It may be the major cause of BF heterogeneity. Large vessel constriction promoted bi-modality in BF histograms. Constriction of large vessels had a greater impact on total transmural BF, as well as transmural BF heterogeneity in comparison to constriction of small vessels. The effect of therapy (adenosine or dialysate cooling) may arise from an increase of total BF, caused by an improved aortic pressure.

CT is now an advanced field of imaging that provides a wide spectrum of BF data in the heart (Cademartiri et al., 2017; Schindler, 2016). In this study, the patient hearts were imaged using a rest/stress protocol. It revealed that interventions have a significant effect. The difference between rest/stress also highlighted inherent vascular structural defects in the heart of patient 3 (Figure 1, second row, last column). Our CT protocols are being continually developed to fully exploit available technologies. However, the BF structure-function appears as a combination of several factors in the imaging, making modelling investigation necessary.

Based on experimental measurements, our model incorporates morphometry data that gave realistic topologies of the coronary vasculature. The optimal assignment of 3D positions to the arterial tree nodes was performed using space filling algorithm (Beard and Bassingthwaighte, 2000). Accounting for limitations, an ensemble of instances was generated whose average haemodynamic properties are presented. The presented models BF heterogeneity (FD = 1.14) is in agreement with FD of 1.2 observed by VanBavel and Spaan in microcirculation SN 2-8 networks (VanBavel and Spaan, 1992). Others have used optimised models, where a spectrum of cost functions such as single arterial volume (Karch et al., 1999; Schreiner et al., 2006), or the combination of metabolic cost, volume, power (Kaimovitz et al., 2005; Keelan et al., 2016) have been optimised. Notwithstanding the relatively simple optimisation used in this study, Figure 4 shows that the BF is distributed throughout most of the myocardium almost uniformly (FD = 1.18). Our model has characteristic transmural BF heterogeneity. Whereas total path lengths of vessels were constrained, the combination of transmural path lengths and the space filling-boundary avoidance algorithm may explain model behaviour. It is known that the myocardium is heterogeneously perfused transmurally (Huo et al., 2009). We therefore computed to alteration of our models transmural heterogeneity as an approximation, and we bear in mind that further development is required.

In this study, multi-modality in BF histograms due to large vessel severe stenosis was observed in the model, which agrees with the imaging data. However, single segment stenosis, or stenosis of high order segments, may be accompanied by an auto-regulatory response. Although overall BF became multi-modal, the FD (computed using the same methods as well as binning) was observed to be reduced. In contrast, the modelling study by Meier et al. (Meier et al., 2004) which included autoregulation showed that FD remained unchanged under stenosis. Remarkably, the inclusion of autoregulation in the above study maintained uni-modality of BF histograms, while increasing RD when low aortic pressure was applied.

The present model was constructed using a limited topology (SN 6 to SN 11). It was also optimised using relatively straightforward space filling and boundary avoidance conditions. Nevertheless, transmural BF heterogeneity was observed, and it was affected by structural alterations. Specifically, blocking the larger (SN 7 or 8) vessels increased transmural heterogeneity more than blocking of the microvasculature (SN 6). Our finding is in line with that of Algranati et al. (Algranati et al., 2011) who have identified sub-endocardial compliance as a cause of BF redistribution. In the above study, they have also identified the contractile factors that may contribute to the sub-endocardial heterogeneity.

Although there have been significant advances in the experimental, theoretical, and imaging literature, we believe that this is one of a few studies that applies coronary vasculature models to indicatively explain clinical imaging observations. Previously developed models have generated 4-D XCAT Phantoms (Fung et al., 2011), or assist the clinician in identifying clinical defects (Fung et al., 2010). Further development of the presented model, at multi-scale and at multi-physics levels, will incorporate Cardiac Physiome models in a pre-clinical assessment tool to assist our clinical research. The model, in conjunction with detailed imaging (see (Jogiya et al., 2014) as well as Figure 1), will expedite the assessment of patient BF status and consequently overall health assessment. The model is also capable of permitting investigation of specific artery bifurcation properties under health and disease conditions (Auricchio et al., 2014). Investigation of properties such as electrical wave propagation and their interaction with vascular structure (Bishop et al., 2010) can be further investigated using the presented model. Extension of the model will incorporate the multi-scale nature of delivery of oxygen to myocardial tissue (Goldman and Zahra Farid, 2017; Mason McClatchey et al., 2017).

## 5. Limitations

### Imaging limitations

Although a large amount of information is available in our clinical images, certain limitations remain. Firstly, only the diastolic phase was collected. During left ventricle’s diastole, the right ventricle is still moving and therefore cannot be imaged. Using a 4D scanning protocol may alleviate this limitation in future studies that can capture right ventricle signals.

Another limitation is that of resolution. Due to the toxicity of the contrast agent and other factors, it is difficult to image the patient for longer durations. However, that results in significantly less number of slices and low resolution. At the acquired resolution, it may not be possible to gain insights into the pre-capillary arteriole BF. Apart from CT, other imaging modalities may be available for assessment of arterial defects (Gharib et al., 2008). The issues regarding toxicity of CT contrast agents may be reduced using magnetic resonance based imaging (Jogiya et al., 2014).

### Modelling limitations

Our model is limited to SN 6 to SN 11 arterial segments. The ensemble of coronary trees generated in this work is based on the morphometric data by the Kassab group. However, the optimisation algorithm in the uneven geometry of the heart is a compute intensive task and it was not possible to generate optimised structures within a short (48 hours) duration. It is known that assigning spatial locations to the vasculature nodes is a large optimisation problem where fractal models based on morphometry need to be combined with theoretical optimisation methods (Schreiner et al., 2006). In future work, we will consider inclusion on further biophysical principles that permit use of cost functions to compute optimal locations of arteriole terminals (Kaimovitz et al., 2010; Keelan et al., 2016; Zamir and Phipps, 1988).

An important simplification of our model is that diameter asymmetry has been ignored. In our model, elements of the same order have a uniform radius. In addition, the bifurcations are assigned randomly where the daughter element radii are assigned according to morphometry rather than according to BF symmetry. Blood vessel diamter asymmetry has been observed in casting data (Huo et al., 2009) and shown to affect BF distributions significantly (Kaimovitz et al., 2008). Bifurcation asymmetry is known to promote increased RD (Sriram et al., 2014) and may be an important factor that contributes to BF heterogeneity in the critically ill groups of patients (Frisbee et al., 2016). It has been quantified in past studies (Dankelman et al., 2007; VanBavel and Spaan, 1992) and will be included in future versions of our model.

Both pulsatile BF (Huo and Kassab, 2006) and viscosity variability in micro-vessels (Pries and Secomb, 2009; Pries et al., 1996) have been ignored for simplicity in this study.

Generation of geometry is based on space filling, whereas optimisation of energy expenditure or some measure will give improved distribution in the future. In the future, we aim to develop a 4D XCT phantom that will permit patient specific BF assessment along with other parameters (Fung et al., 2011).

The lack of autoregulation is a significant simplification in our study. The inclusion of autoregulation may affect results presented in this study (Meier et al., 2004). It will also permit testing of further factors that affect local and global flow, as shown recently by Namani et al. (Namani et al., 2017).

Apart from structural changes, BF is also affected by several other factors, where metabolic demand of tissue and protein expression heterogeneity are prominent (Stoll et al., 2008). Autoregulation, as well as other patho-physiological factors will be incorporated into the model in future work.

## 6. Clinical significance of the study

In this study, imaging data acquired from patients was analysed. Whereas it is probably true that dialysis causes vascular dysfunction by affecting the micro-vasculature, this study’s imaging observation combined with the modelling results, indicates that large vessel dysfunction may significantly affect patient’s coronary perfusion.

In the process of quantifying BF heterogeneity, we have now developed algorithms that compute simple yet informative measures of BF heterogeneity. Such a tool will provide rapid assessment of whether imaging data reflect the effectiveness of therapy.

An important development during this study was the implementation of a method to generate organ level vascular structure. When combined with micro-vasculature, the vasculature model is being developed to become a pre-clinical trial in silico indicative outcome assessment tool.

**Author Contributions** SRK, DG, and CWM designed the study in consultation with CE. AS and TL provided the texture analysed CT data and related text. SRK and FS segmented the hearts. SRK developed the codes and performed the simulation experiments, acquired, curated, and analysed the data. DG and CWM provided expert comments. CE provided expert insights into clinical-modelling data interpretation. SRK wrote the first draft. AS provided the draft of the imaging data analysis. All authors wrote and approved the final manuscript.

## Acknowledgements

We thank LHSC IT, Compute Canada, SHARCNet for IT and HPC resources. This study was supported by Heart and Stroke grant (G-17-0018311, PI: CWM).

